# Novel mouse model of encephalocele: post-neurulation origin and relationship to open neural tube defects

**DOI:** 10.1101/649624

**Authors:** Ana Rolo, Gabriel L. Galea, Dawn Savery, Nicholas D. E. Greene, Andrew J. Copp

**Affiliations:** Newlife Birth Defects Research Centre, GOS Institute of Child Health, University College London, 30 Guilford Street, London WC1N 1EH, UK

## Abstract

Encephalocele is a clinically important birth defect that can lead to severe disability in childhood and beyond. The embryonic pathogenesis of encephalocele is poorly understood and, while usually classified as a ‘neural tube defect’, there is conflicting evidence on whether encephalocele results from defective neural tube closure, or is a post-neurulation defect. It is also unclear whether encephalocele can result from the same causative factors as anencephaly and open spina bifida, or whether it is aetiologically distinct. This lack of information results largely from the scarce availability of animal models of encephalocele, particularly ones that resemble the commonest, non-syndromic human defects. Here we report a novel mouse model of occipito-parietal encephalocele, in which the small GTPase Rac1 is conditionally ablated in the (non-neural) surface ectoderm. Most mutant fetuses have open spina bifida, and some also exhibit exencephaly/anencephaly. However, a large proportion of mutant fetuses exhibit encephalocele affecting the occipito-parietal region. The encephalocele phenotype does not result from a defect in neural tube closure, but rather from a later disruption of the surface ectoderm covering the already closed neural tube, allowing the brain to herniate. The neuroepithelium itself shows no down-regulation of Rac1 and appears morphologically normal until late gestation. A large skull defect develops overlying the region of brain herniation. Our work provides a new genetic model of occipito-parietal encephalocele, particularly resembling non-syndromic human cases. While encephalocele has a different, later-arising pathogenesis than open neural tube defects, both can share the same genetic causation.

**SUMMARY STATEMENT:** Encephalocele - a severe brain defect - arises after neural tube closure, but can share a common genetic cause with anencephaly, a defect of neural tube closure.

## INTRODUCTION

Encephalocele is a severe birth defect of the skull and brain with a median prevalence of 0.1-0.3 per 1000 births, although with geographical variation (1). The meninges, with/without brain tissue, herniate outside the skull as a sac, exposing the brain to potential damage both pre- and post-natally. Despite surgical repair soon after birth, later health problems are common, including hydrocephalus, epilepsy and learning difficulties. Encephaloceles emerge along the skull midline, with variation in rostro-caudal location which can be fronto-ethmoidal, parietal, occipital or cervical. Generally, the prognosis worsens with posterior location, size of sac and increasing amount of herniated brain tissue (2).

Although most cases are sporadic and of unknown causation, encephalocele can form part of a syndrome as in trisomy 18, Knoblock syndrome (*COL18A1* mutation), amniotic band syndrome and warfarin embryopathy (3). Occipital encephalocele is best known as part of Meckel syndrome (overlapping with Joubert syndrome), in which individuals also exhibit polydactyly, polycystic kidneys and biliary defects. In recent years, mutations in several genes (e.g. *MKS1, MKS2* (*TMEM216), MKS3* (*TMEM67), CEP290, RPGRIP1L*) have been identified in various forms of Meckel syndrome (4). Cellular analysis of the MKS proteins has demonstrated a key role in the structure and function of primary cilia, and Meckel syndrome is thus now classified as a ciliopathy.

Mice that incorporate mutations of some of the genes responsible for Meckel syndrome display biliary, limb and kidney defects resembling the human syndrome, as well as defective ciliary structure and/or function (5-7). While failure of cranial neural tube closure was described in a proportion of *Tmem67* (MKS3) null mice (8), none of the mouse models appear to exhibit herniation of brain tissue outside the skull, which would represent an encephalocele.

While often classified as a ‘neural tube defect’ (NTD) (4;9), the embryonic/fetal pathogenesis of encephalocele is less well understood than for other NTDs, particularly anencephaly and open spina bifida. The latter conditions result from defective closure of the neural tube (i.e. primary neurulation), as demonstrated by studies of NTD pathogenesis in mouse mutants (10). Of the many (> 240) mouse mutants so far described, very few display a phenotype corresponding to encephalocele (11). Hence, the mouse data do not yet conclusively shed light on whether encephalocele is a primary neurulation defect or a post-neurulation anomaly, such as herniation of the closed neural tube through a skull defect.

Recently, encephalocele was identified as part of a ‘cluster’ of NTDs in the pregnancies of HIV-positive Botswanan women exposed to the anti-retroviral drug dolutegravir from the time of conception (12). NTDs in this cluster comprised one case each of encephalocele, anencephaly, myelomeningocele and iniencephaly, and occurred with a prevalence of 9.4 per 1000 births (4 out of 426 infants). This is a markedly elevated prevalence, compared with 1.2 per 1000 or less in comparator groups that were not exposed to dolutegravir from conception. An urgent need is to determine whether this cluster of NTDs represents a causal association with dolutegravir. However, this is hampered by uncertainty over whether encephalocele can share causation with ‘open’ NTDs: anencephaly and myelomeningocele.

Here we describe a mouse model of encephalocele resulting from conditional deletion of *Rac1*, a small GTPase of the Rho family, in the non-neural (surface) ectoderm of the embryo. These mice exhibit open spina bifida (myelomeningocele equivalent) and, in some cases exencephaly (anencephaly equivalent) (13-15). We show that a large proportion of these mice also develop occipito-parietal encephalocele, detectable from embryonic day (E) 13.5 onwards. The encephalocele displays a fully closed neural tube at the level of the lesion, with an associated overlying skull defect. Hence, encephalocele is a post-neurulation anomaly that can be caused by the same genetic defect as open spina bifida and exencephaly/anencephaly.

## RESULTS

### Grhl3Cre-Rac1 mutants display exencephaly or encephalocele

*Rac1* was conditionally deleted by expressing Cre recombinase under control of the *Grhl3* promoter. *Grhl3*^*Cre/+*^; *Rac1*^*f/-*^ and *Grhl3*^*Cre/+*^; *Rac1*^*f/f*^ genotypes both lack Rac1 expression mainly in the surface ectoderm (14), and do not differ morphologically. Hence, these genotypes were pooled for analysis and are denoted ‘Grhl3Cre-Rac1’. They were compared with Cre-expressing control littermates: *Grhl3*^*Cre/+*^; *Rac1*^*f/+*^ and *Grhl3*^*Cre/+*^; *Rac1*^*+/-*^, which retain Rac1 expression (denoted ‘Grhl3Cre-Con’; Figure 1A). Littermates without Cre expression (*Grhl3*^*+/+*^; *Rac1*^*f/f, f/+, f/-or +/-*^) were denoted ‘Non-Cre’ controls (Table 1).

**Table 1.**
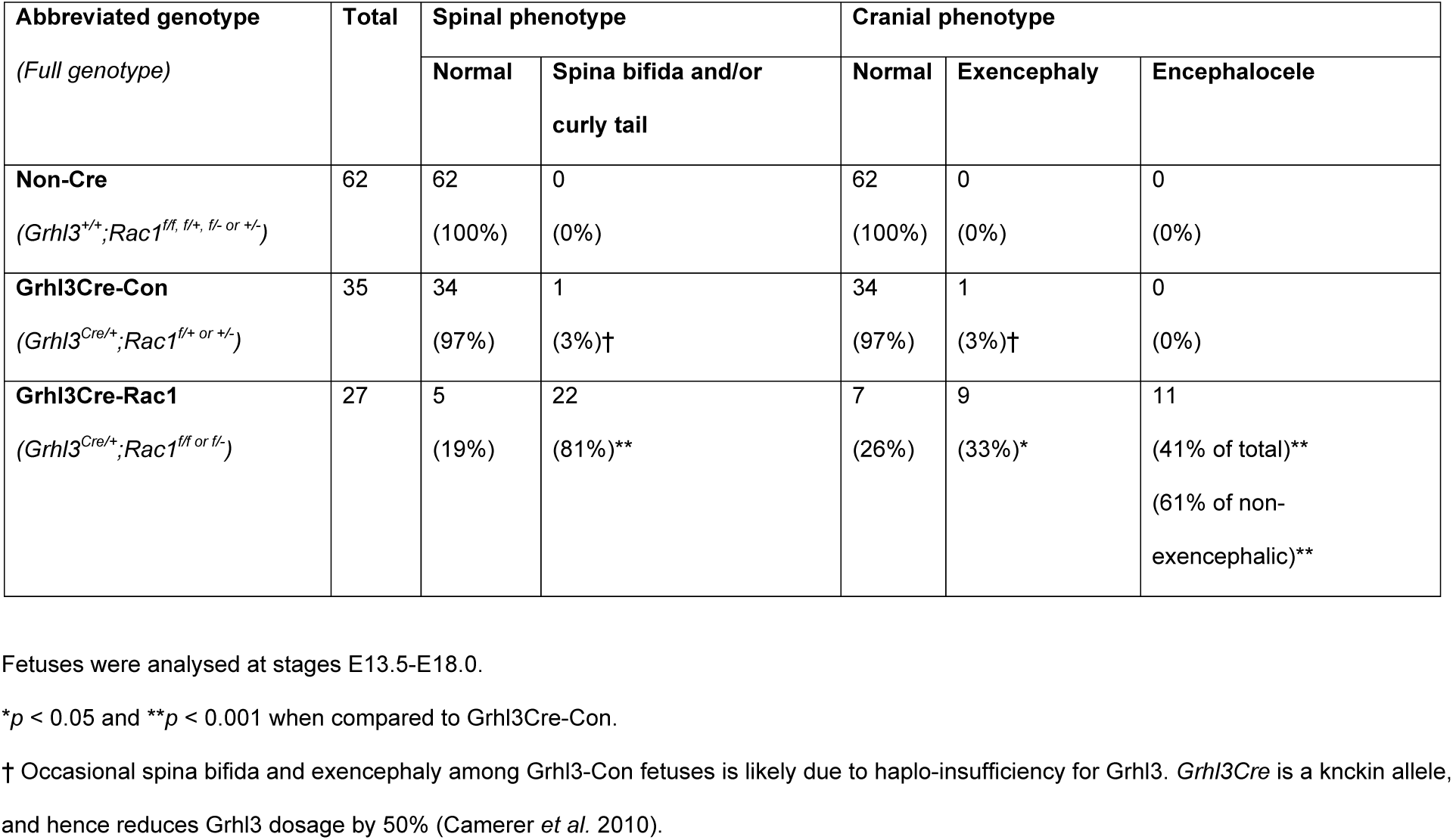
Genetic analysis of Grhl3Cre-Rac1 conditional mutants

| Abbreviated genotype<br>(Full genotype) | Total | Spinal phenotype |  | Cranial phenotype |  |  |
| --- | --- | --- | --- | --- | --- | --- |
|  |  | Normal | Spina bifida and/or curly tail | Normal | Exencephaly | Encephalocele |
| **Non-Cre**<br>(*Grhl3*<sup>+/+</sup>; *Rac1*<sup>f/f, f/+, f/- or +/-</sup>) | 62 | 62<br>(100%) | 0<br>(0%) | 62<br>(100%) | 0<br>(0%) | 0<br>(0%) |
| **Grhl3Cre-Con**<br>(*Grhl3*<sup>Cre/+</sup>; *Rac1*<sup>f/+ or +/-</sup>) | 35 | 34<br>(97%) | 1<br>(3%)† | 34<br>(97%) | 1<br>(3%)† | 0<br>(0%) |
| **Grhl3Cre-Rac1**<br>(*Grhl3*<sup>Cre/+</sup>; *Rac1*<sup>f/f or f/-</sup>) | 27 | 5<br>(19%) | 22<br>(81%)** | 7<br>(26%) | 9<br>(33%)* | 11<br>(41% of total)**<br>(61% of non-exencephalic)** |

Fetuses were analysed at stages E13.5-E18.0.
\* $p < 0.05$ and \*\* $p < 0.001$ when compared to Grhl3Cre-Con.
† Occasional spina bifida and exencephaly among Grhl3-Con fetuses is likely due to haplo-insufficiency for Grhl3. *Grhl3Cre* is a knckin allele, and hence reduces Grhl3 dosage by 50% (Camerer *et al.* 2010).

**Figure 1.**
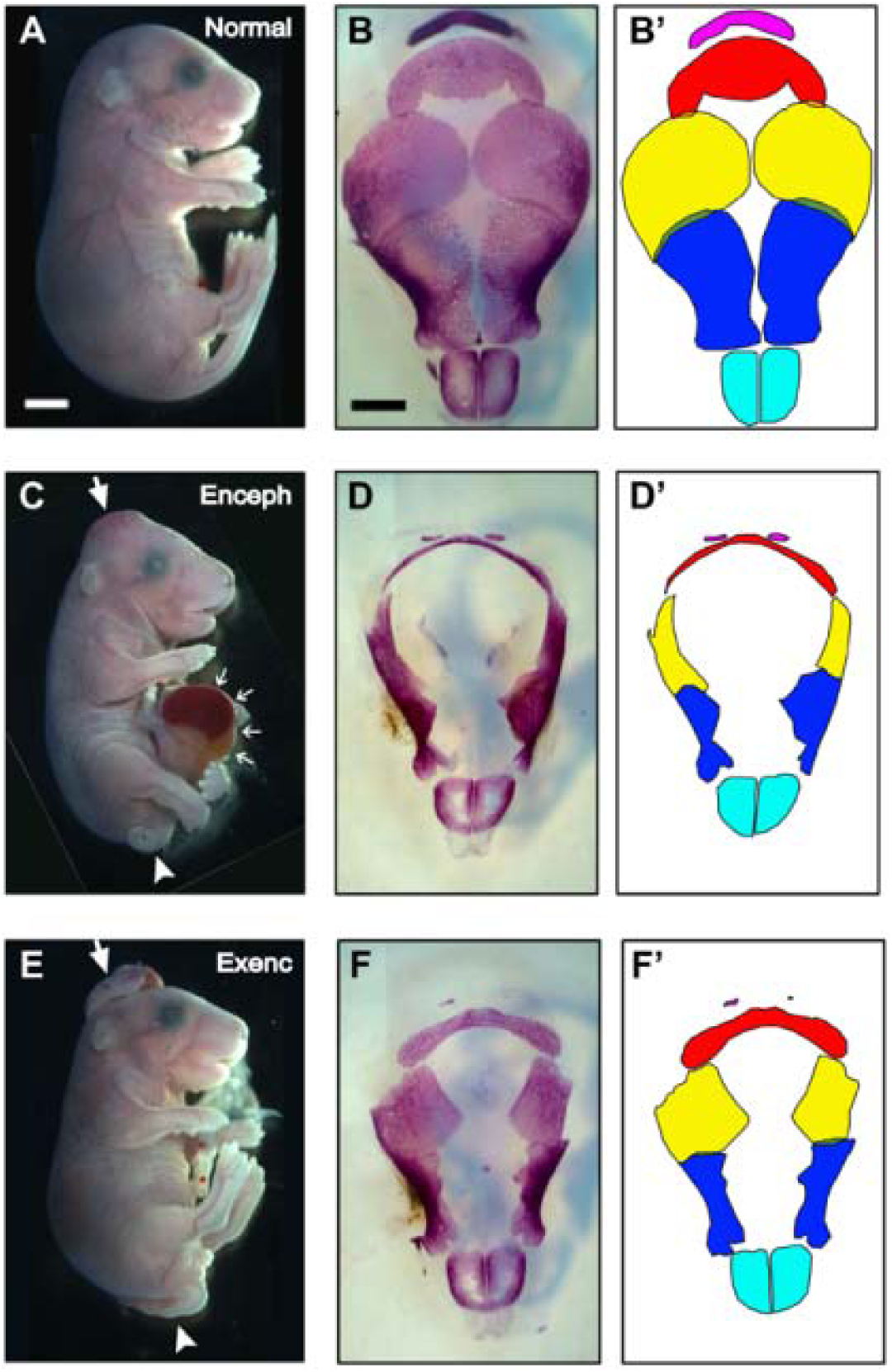
Grhl3Cre-Rac1 mutants display exencephaly or encephalocele, both accompanied by skull defects. Mice shown are: (A,B,B’) Grhl3Cre-Con; (C,D,D’) and (E,F,F’) Grhl3Cre-Rac1 conditional mutants at E17.5. Note the encephalocele brain protrusion in a mutant fetus (arrow in C) and the open NTD exencephaly in another mutant fetus (arrow in E). Both mutants have tail defects (arrowheads in C, E). The fetus with encephalocele also has a ventral body wall defect, exposing internal organs (small arrows in C). This was seen in 3/8 mutant fetuses (not specifically associated with encephalocele) from stage E16.5 onwards, and in none of 17 controls. (B, D, F) Skull preparations (calvaria only shown, from top view). (B’, D’, F’) Diagrams show the identity of the bones (top to bottom): pink, occipital; red, inter-parietal; yellow, parietal; blue, frontal; cyan, nasal. The bones of the Grhl3Cre-Con fetus are well formed and meet in the dorsal midline, prefiguring the sagittal suture. In contrast, a large midline deficit in bone formation occurs in both mutant fetuses. In the fetus with encephalocele, all bones except the nasals are severely affected in their medial aspects (D, D’). Similar, but less severe, defects are present in the fetus with exencephaly (F, F’). Analyses performed on at least 3 different embryos of each genotype and phenotype group; representative specimens are shown. Scale bars: 1 mm (A, C, E) and 500 µm (B, D, F).

Grhl3Cre-Rac1 mutants develop spina bifida at high penetrance (81%; Figure 1C, E; Table 1), as described previously (13). Exencephaly (Figure 1E) also occurs in mutants, but at lower frequency affecting 30% (21/69) of Grhl3Cre-Rac1 embryos at E9.5 and 25% (11/44) at E10.5-13.5 (14). Here we examined E13.5-17.5 fetuses and found exencephaly in 33% (9/27; Table 1). Hence, exencephaly is first seen at the stage when cranial neural tube closure is usually completed (E9.5) and persists at a relatively constant rate into later gestation.

From E13.5 onwards, we also encountered another cranial phenotype, resembling parieto-occipital encephalocele (Figure 1C). This affected more than half (61%) of the non-exencephalic Grhl3Cre-Rac1 fetuses (41% of total fetuses) and was not seen in littermate controls (Table 1; Figure 1A). Encephalocele appeared as a smooth protrusion of the cranial region, in contrast to the irregular ‘mushroom-like’ appearance of exencephaly (compare Figure 1C and E).

Hence, both exencephaly and encephalocele occur among mouse mutants lacking Rac1 expression in the surface ectoderm, but the two defects arise in different individuals, at different developmental stages: exencephaly at neurulation, and encephalocele post-neurulation.

### Larger skull defects in mutants with encephalocele than exencephaly

Skull preparations showed that at E17.5 the bones of Grhl3Cre-Con fetuses are well formed and meet at the dorsal midline, prefiguring the sagittal suture (Figure 1B, B’). In contrast, Grhl3Cre-Rac1 mutant fetuses with encephalocele had a large midline deficit in bone formation, where all bones except the nasals were severely affected in their medial aspects (Figure 1 D, D’). Similar bone defects were seen in fetuses with exencephaly (Figure 1 F, F’), but strikingly these were less severe, despite the very pronounced exencephalic brain defect (Figure 1E).

Histological sections through the affected regions of Grhl3Cre-Rac1 mutants with encephalocele at E17.5 showed that the encephalocele protrusion, besides lacking a bony tissue covering, was also devoid of overlying epidermal and mesenchymal tissues (compare Figure 2C, F). In contrast, sections through head regions rostral to the encephalocele displayed brain covered by epidermal tissue and mesenchyme (compare Figure 2B, E), despite the lack of bone, as expected from the skull preparation analyses.

**Figure 2.**
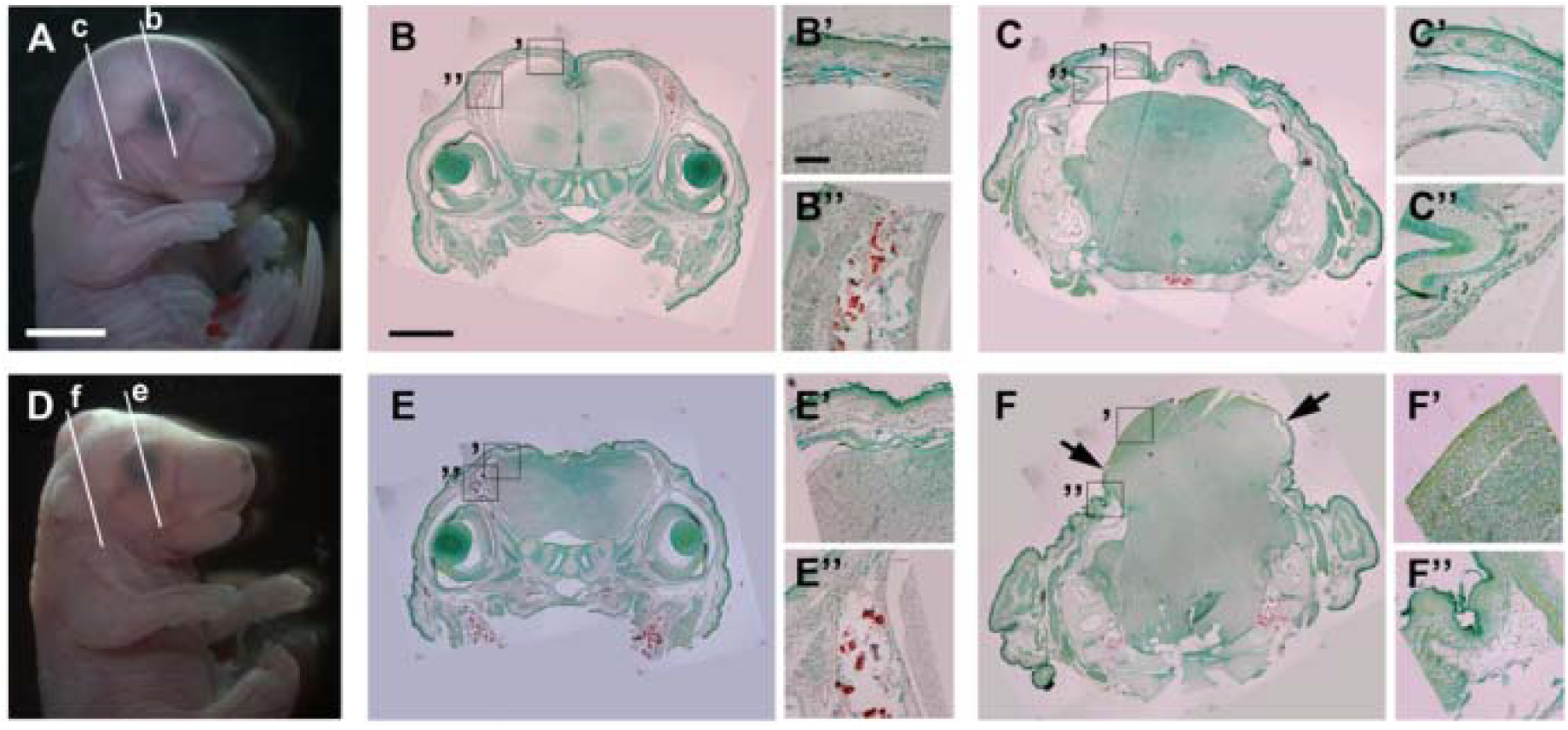
Encephalocele in Grhl3Cre-Rac1 mutants is not covered by skin or bone tissue. Mice shown are: (A-C) Grhl3Cre-Con; (D-F) Grhl3Cre-Rac1 conditional mutants at E17.5. Low magnification sections through the head (B, C, E, F), stained with Alizarin Red to reveal mineralised bone, are taken from the levels indicated in A and D. Boxed areas in the low magnification sections are shown at higher magnification in B’, B”, C’, C”, E’, E”, F’, F”. Note the encephalocele brain protrusion in the Grhl3Cre-Rac1 fetus (D). The section taken rostral to the encephalocele (E; at level of the eyes) shows a closed neural tube covered by epidermis. However, the dorsal surface is irregular compared with the smooth appearance of the Grhl3Cre-Con fetus (B). The section taken through the encephalocele (F) shows a massive extrusion of brain tissue from the dorsal surface of the head. Epidermis and subcutaneous tissue are present at the edge of the encephalocele (F”) but do not cover the brain protrusion (F’), in contrast to the non-mutant appearance (C’). The brain appears ‘closed’ (i.e. normally neurulated) in the protruded area, although there is clear internal disorganisation. Analyses performed on at least 3 different embryos of each genotype and phenotype group; representative specimens are shown. Scale bars: 2 mm (A, D), 500 µm (B, C, E, F) and 100 µm (B’, B”, C’, C”, E’, E”, F’, F”).

### Encephalocele development is preceded by a defect in the surface ectoderm

The brain tissue in Grhl3Cre-Rac1 E17.5 mutants with encephalocele appeared ‘closed’ (i.e., normally neurulated), despite a disrupted internal organization (Figure 2E, F). To further address the developmental origins of the encephalocele, we examined sections through the mid- and hind-brain of earlier stage embryos, at E13.5, when the encephalocele defect first becomes identifiable. At this stage the brain tissue was still covered by surface ectoderm and mesenchymal layers (Figure 3B, C, E, F), although breaks and discontinuities could already be observed in these layers (Figure 3E”, F”, arrows). The neural tissue, however, was morphologically similar to that in control embryos, with a completely closed neural tube and a well-defined ventricular lumen at all rostro-caudal levels of the brain, including at the levels where the overlying tissues appeared broken. This finding not only confirmed that the encephalocele does not result from a failure of neural tube closure, but also showed that the brain defects observed at E17.5 develop after the onset of encephalocele.

**Figure 3.**
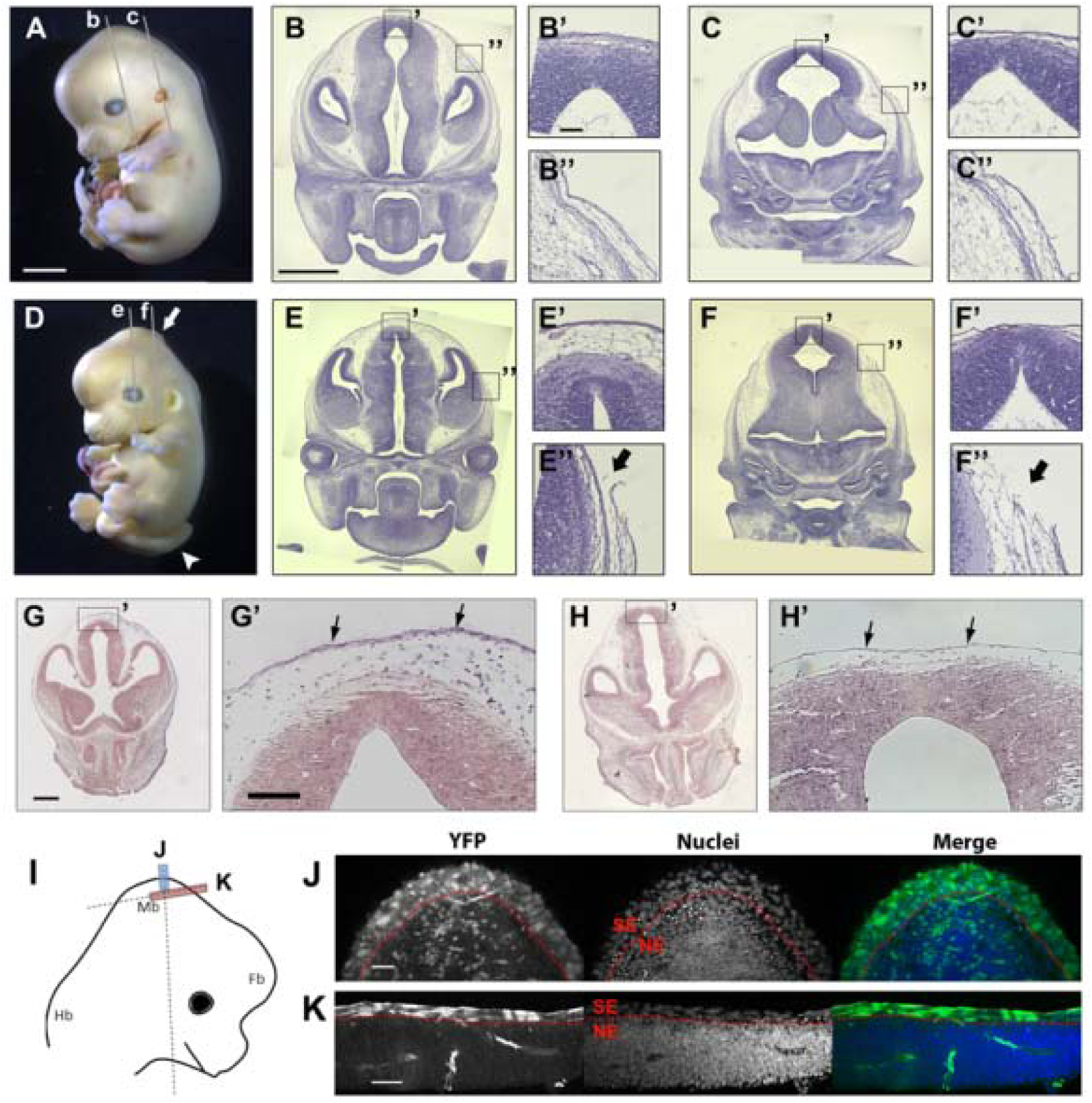
First appearance of encephalocele in Grhl3Cre-Rac1 mutants. Mice shown are: (A-C, G) E13.5 Grhl3Cre-Con; (D-F, H) E13.5 Grhl3Cre-Rac1 conditional mutant. Low magnification sections through the head at E13.5 (B, C, E, F), stained with H&E, are taken from the levels indicated in A and D. Boxed areas in the low magnification sections are shown at higher magnification in B’, B”, C’, C”, E’, E”, F’, F”. Note the encephalocele brain protrusions (arrow in D) in the mutant fetus, which also exhibits open spina bifida (white arrowhead in D). Disruption of the epidermis and subcutaneous tissue at the edge of the brain protrusions is indicated by black arrows (in E”, F”). The comparable tissues in the non-mutant fetus are intact (B”, C”). In situ hybridisation in coronal sections through the midbrain region at E12.5, using an antisense RNA probe against mouse Rac1 exons 4 and 5 show Rac1 mRNA is expressed (purple staining) in all tissues of Grhl3Cre-Con (G, G’) and in neural tube of the Grhl3Cre-Rac1 conditional mutant (H, H’). Expression is specifically abolished in the mutant surface ectoderm (black arrows in H’) where Grhl3Cre causes recombination of the floxed Rac1 alleles. At this stage, the mutant shows a completely closed neural tube, covered by intact surface ectoderm, similar to control. (I) Schematic of an E12.5 mouse embryo head with dashed lines indicating the planes of section shown in J,K. Coloured boxes show the areas and planes of section. (J, K) Grhl3Cre-driven recombination of the ROSA26-YFP-reporter, as detected by anti-YFP immunofluorescence. Cre-mediated recombination can be seen in all surface ectoderm cells (above dotted red line), but only in a few scattered neuroepithelial cells (below dotted red line). Analyses performed on at least 3 different embryos of each genotype and phenotype group; representative specimens are shown. Scale bars: 500 µm (A, D), 100 µm (B, C, E, F), 20 µm (B’, B”, C’, C”, E’, E”, F’, F”), 100 µm (G, H), 50 µm (G’, H’, J, K).

To more precisely pinpoint the timing of encephalocele development, we analysed Grhl3Cre-Rac1 mutant embryos at E12.5 (Figure 3G, H). Use of the Rosa26-EYFP reporter confirmed that, as at earlier stages (Rolo et al, 2016), Grhl3Cre-driven recombination at E12.5 is present in the entire dorsal surface ectoderm, but only in a variable small minority of cells in the neuroepithelium and mesenchyme (Figure S1). Despite Rac1 knock-down in all surface ectoderm cells (Figure 3H’), the tissues overlying the brain of mutant embryos were still intact at E12.5 and morphologically similar, although apparently thinner, compared with those of Grhl3Cre-Con embryos (Figure 3G, H). We conclude that encephalocele in Grhl3Cre-Rac1 mice arises between E12.5 and E13.5, following completion of cranial neural tube closure, when the surface ectoderm and associated tissues are lost and the neural tube becomes exposed to the external environment.

## DISCUSSION

Over 240 mouse models of NTDs have been reported, mainly displaying exencephaly, spina bifida, or both (11). To date, however, only two convincing mouse models of encephalocele have been described: the *tuft* mouse which involves a mutation of the *Tet1* gene (20;21) and the *fog* mutant in which the *Apaf1* gene is mutated (22). Both display frontal encephalocele together with craniofacial defects and, in *tuft* mice, also lipoma. In humans, fronto-ethmoidal encephalocele is particularly found in South-East Asia (23), whereas in other geographical locations it is a less common defect than occipital encephalocele. Moreover, lipoma does not typically accompany any of the varieties of human encephalocele. The defect we describe here, in Grhl3Cre-Rac1 mutants, is the first report to our knowledge of a murine model of occipito-parietal encephalocele, and without accompanying facial defects or lipoma. It therefore represents a ‘proof-of-principle’ study of the origin during brain development of a type of encephalocele that is common in humans.

Even though encephaloceles are increasingly classified as post-neurulation defects, uncertainty continues over their relationship to neural tube closure. Indeed, the encephalocele in *tuft* mice has been described as resulting from incomplete closure of the anterior neural tube (20). On the other hand, some authorities argue that encephalocele is a later-arising defect, resulting from incomplete fusion of the skull bones at the midline, creating a gap through which meninges and brain tissue herniate (23). The defect in Grhl3Cre-Rac1 mutants is first detected at E13.5, approximately 4 days after anterior neural tube closure is completed, but before the beginning of skull ossification. In sharp contrast, exencephaly arises in the same mutant litters, with an onset at the stage of cranial neural tube closure.

We conclude, therefore, that encephalocele in Grhl3Cre-Rac1 mutants is neither a result of a failure in neurulation nor primarily a skull defect. Rather, it develops after neural tube closure due to a defect in the surface ectoderm, and the defect is already manifest by the time of skull formation. Despite not being the result of a skull defect, encephalocele in Grhl3Cre-Rac1 mutants is nonetheless associated with severe malformation of skull bone formation, likely explaining the later pathogenesis of human encephalocele, in which the brain and/or meninges herniate through a skull defect.

Besides the occipito-parietal location of the encephalocele, another difference between our study and the *tuft* and *fog* mouse models of frontal encephalocele, is the lack of skin coverage in Grhl3Cre-Rac1 mutants. In humans, the encephalocele sac is often skin-covered but this is not universal. For example, in one series (24), only 2 out of 12 large encephaloceles had skin coverage. Whether absence of skin is a primary feature, or a secondary degenerative change, is not clear. Indeed, loss of skin over encephaloceles has been documented (25). Surface damage as a result of exposure to amniotic fluid is possible, leading to loss of epidermis and potentially damage to herniated brain tissue. Degeneration of neural tissue exposed to amniotic fluid was demonstrated most clearly in the transition from exencephaly to anencephaly (26), and is part of the rationale for introducing surgical closure of open spina bifida during the fetal period, as an alternative to postnatal surgery (27).

Encephalocele occurs in humans both sporadically, accounting for 50-75% of cases (9;28), and syndromically as typified by Meckel syndrome. While the genetic basis of Meckel syndrome, as a ciliopathy, has been established in recent years (4), the developmental link between ciliopathy and encephalocele in this condition remains unclear. Our finding of tissue-specific Rac1 deletion causing mouse occipito-parietal encephalocele may relate most closely to the origins of human non-syndromic encephalocele, where causation and pathogenesis are least understood.

Rac1 is required for many cellular processes including maintenance of cell proliferation, integrity of epithelial cell junctions and cytoskeletal events in cell shape change and motility (29) Constitutive inactivation of Rac1 is lethal at an early embryonic stage, before neurulation begins (30), and so Rac1 function in vivo has been investigated by conditional gene targeting studies, as in the present study. For example, tissue-specific depletion of Rac1 in the early embryo causes defective cell migration, both of the anterior visceral endoderm (31) which is required for head induction, and of the mesoderm during subsequent gastrulation (32). Neural crest migration and differentiation are defective in the absence of Rac1 (33-35). Both canonical and non-canonical Wnt signalling require Rac1 for full function (36;37). Most important for the present study is the finding that inactivation of Rac1 in adult skin led epidermal stem cells to exit from the cell cycle and undergo differentiation (38). Hence, loss of Rac1 in the surface ectoderm overlying the brain in our study likely induces faulty tissue expansion and hence disruption of tissue integrity, enabling brain herniation. We conclude that Rac1 and the several signalling pathways in which it functions may provide a focus for future studies of the genomic and/or epigenomic changes that predispose to non-syndromic encephalocele.

The embryonic pathogenesis of encephalocele, as a post-neurulation defect, shows it to be quite distinct from the neural tube closure defects: anencephaly and myelomeningocele. Nevertheless, an important finding of the present study is that both encephalocele and exencephaly/anencephaly can result from the same genetic insult: loss of Rac1 in the surface ectoderm. It is not yet clear why some individuals fail in cranial neural tube closure and yet other genetically identical individuals complete closure but subsequently develop brain herniation. An important implication of our findings is, however, that in cases of suspected environmental exposures, as in HIV-positive women exposed during early pregnancy to the antiretroviral drug dolutegravir (12), it is perfectly plausible that cases of encephalocele, anencephaly and myelomeningocele could all be etiologically related, as alternative outcomes of faulty nervous system development in the embryo.

## MATERIALS AND METHODS

### Mouse procedures and experimental design

Mouse studies were conducted under the auspices of the UK Animals (Scientific Procedures) Act 1986, as described in Project Licence 70-7469 which was scrutinised and approved by the Animal Welfare and Ethical Review Body of University College London. Mice were housed under standard conditions with environmental enrichment. Strains were: *Grhl3*^*Cre/+*^ (13), *Rac1*^*f/f*^ (16), and *ROSA26-EYFP* (17), all on a C57BL/6 background. Matings were: *Grhl3*^*Cre/+*^; *Rac1*^*f/+ or -/+*^ x *Rac1*^*f/f*^ (14). Embryos were dissected in Dulbecco’s modified Eagle’s Medium (DMEM; Invitrogen) containing 10% fetal bovine serum (Sigma), and rinsed in phosphate buffered saline (PBS) prior to fixation. Embryo genotyping was by PCR of yolk sac DNA, as described (14). Experiments were conducted according to the ARRIVE guidelines (www.nc3rs.org.uk): e.g. all analyses were performed blind to genotype which was obtained after data collection had been completed.

### Histology and skull preparations

Embryos were fixed over several days in Bouin’s solution (Sigma) or in 4% paraformaldehyde in PBS, dehydrated in an ethanol series and embedded in paraffin-wax. Sections (5 µm thickness) were stained with Harris’ haematoxylin solution and 2% Eosin Y, or Alizarin Red and Fast Green (all Sigma). Images were captured on an Axiophot2 upright microscope. For skull preparations, embryo heads were skinned and stained with Alizarin Red (0.15%) in 1% KOH and cleared with 1% KOH in 20% glycerol (18).

### mRNA in situ hybridisation

In situ hybridisation was performed on 5 µm-thick paraffin sections using a digoxigenin-labelled anti-sense RNA probe designed to detect the exons deleted in the *Rac1* conditional mutant (14). Images were captured on an Axiophot2 upright microscope.

### Immunofluorescence

Embryos were fixed for 24 h in 4% paraformaldehyde in PBS, pH 7.4, at 4°C. Immuno-fluorescence for YFP (yellow fluorescent protein) was performed on 12 µm-thick cryosections of gelatine-embedded embryos (7.5% gelatine [Sigma] in 15% sucrose) using an anti-GFP (green fluorescent protein) rabbit polyclonal Alexa488-conjugated antibody (Life Technologies A21311) at 1:400 dilution. Anti-GFP cross-reacts with YFP. Nuclei were labelled with TO-PRO-3 (Thermo Fisher). Images were captured on an LSM880 Examiner confocal system (Carl Zeiss Ltd, UK) as previously reported (19), and linear adjustments made using Fiji software.

### Statistics

Fisher’s exact test was used for comparison of phenotype frequencies in Table 1. When more than two groups were compared in multiple tests, the chance of detecting false positives was minimised by ‘protecting the alpha-level’: i.e. the conventional *p* = 0.05 critical value was divided by the number of tests. Hence, for 2 tests, *p* < 0.025 was interpreted as indicating *p* < 0.05.

## ACKNOWLEDGMENTS

We thank Berta Crespo for assistance with in situ hybridisation. This research was supported by programme grant 087525 from the Wellcome Trust.

